# Eye movements during text reading align with the rate of speech production

**DOI:** 10.1101/391896

**Authors:** Benjamin Gagl, Klara Gregorova, Julius Golch, Stefan Hawelka, Jona Sassenhagen, Alessandro Tavano, David Poeppel, Christian J. Fiebach

## Abstract

Across languages, the speech signal is characterized by a predominant modulation of the amplitude spectrum between about 4.3-5.5Hz, reflecting the production and processing of linguistic information chunks (syllables, words) every ∼200ms. Interestingly, ∼200ms is also the typical duration of eye fixations during reading. Prompted by this observation, we demonstrate that German readers sample written text at ∼5Hz. A subsequent meta-analysis with 142 studies from 14 languages replicates this result, but also shows that sampling frequencies vary across languages between 3.9Hz and 5.2Hz, and that this variation systematically depends on the complexity of the writing systems (character-based vs. alphabetic systems, orthographic transparency). Finally, we demonstrate empirically a positive correlation between speech spectrum and eye-movement sampling in low-skilled readers. Based on this convergent evidence, we propose that during reading, our brain’s linguistic processing systems imprint a preferred processing rate, i.e., the rate of spoken language production and perception, onto the oculomotor system.

---

Speech production and perception form a quasi-rhythmic information processing cycle^1^. During spoken communication, our brain entrains to the frequency structure of the speech signal^2,3^, suggesting that the temporal structure of the linguistic stimulus drives neural processes in auditory and language processing systems^4^. Across languages, the amplitude modulation spectrum of natural speech peaks consistently in a frequency range between 4.3 and 5.5Hz^5,6^, which reflects that informative signals (e.g., syllables^7,8^) are processed by the listeners’ brains every ∼200ms^9^. Interestingly – and we hypothesize not accidentally – a typical eye fixation during reading has a very similar duration, i.e., between ∼200ms for orthographically transparent writing systems like German or Finnish^10,11^ and ∼250ms for character-based systems like Chinese^11,12^.

Abundant research has used eye-movement recordings to study reading at high temporal resolution, exploring, for example, how reading is influenced by word length, word frequency, or word predictability given a sentence context^12^. Among various measures that can be derived from eye-movement recordings, timing measures like fixation duration are most frequently examined and considered precise markers of reading speed^13,14^. These temporally highly-resolved measurements have so far only been analyzed at the level of individual items – typically words. However, other domains of cognitive research (like attention^15^) demonstrate that eye movements can also be subjected to frequency-based analyses. We here demonstrate that a frequency-based exploration of how written text is sampled by the eyes can open up new perspectives onto several fundamental questions related to the process of reading, including whether reading is related to spoken language processing as recent investigations of word-per-minute measures suggest^16^ and whether the visual system’s sampling of linguistic input differs from eye movements during non-linguistic tasks or between different languages or writing systems.

To address these foundational questions, we first used an empirical dataset^17^ to determine eye-movement sampling frequencies for 50 native speakers of German during sentence reading compared to a non-linguistic control task, using two different methodologies. Next, to determine the generality of these results and to investigate possible cross-linguistic differences in the sampling rate of reading, we conducted a meta-analysis of 124 studies from 14 different languages. To this end, we established a frequency analysis for fixation durations extracted from published eye-tracking studies. Finally, we acquired two novel datasets, one with 48 non-native and one with 86 native speakers of German, to investigate directly the relationship between the sampling frequency of reading and speech production rates on a subject-by-subject level. Experimental and meta-analytic results show (i) that written text is sampled in the same frequency range as spoken language, (ii) that the sampling rate of reading has an upper limit at ∼5Hz, observable in languages with transparent orthographies, (iii) that this rate can be modulated depending on the complexity of the writing system (e.g., in character-based as opposed to alphabetic scripts or in alphabetic scripts with opaque grapheme-phoneme mapping), and (iv) that a direct coupling between reading and speech rates is only found in persons with lower levels of reading skill.

## Results: Estimating the sampling rate of reading

50 healthy volunteers read sentences from the Potsdam Sentence Corpus (144 sentences presented as a whole; 1,138 words in total; see Ref.^10^) while movements of their right eye were tracked (resolution: 1,000Hz). As non-linguistic control task, participants scanned ‘z-strings’ that were constructed by replacing all letters of the sentence by the letter ‘z’ (e.g., “*Ein berühmter Maler hat sich selbst ein Ohr abgeschnitten”/ A famous painter cut off his own ear*. was transformed to “*Zzz zzzzzzzzz Zzzzz zzz zzzz zzzzzz zzz Zzz zzzzzzzzzzzzz*.“; see Methods for details and Ref.^17^ for previous results from this dataset). Given that fixation numbers do not differ significantly between sentences and z-strings^17–19^, similar scan paths are assumed which qualifies z-strings as valid control stimuli for reading experiments (see Supplementary Information 1 for a detailed comparison of scan-paths).

### Fixation durations

After preprocessing (leading to removal of 3.1% of the data), we estimated mean fixation durations separately for each participant and experimental condition. Figure 1a shows that fixation durations (presented here as subject-specific means) are shorter for reading than scanning (average: 197ms vs. 249ms, respectively; Effect size: 52ms; Cohen’s d =1.57; *t*(49)=11.1; p<.001). This has been reported previously for this dataset^17^ and replicates earlier results for German^18^, English^20^, and French^19^ in which fixation durations increased from reading to scanning between 38-42ms.

**Figure 1.**
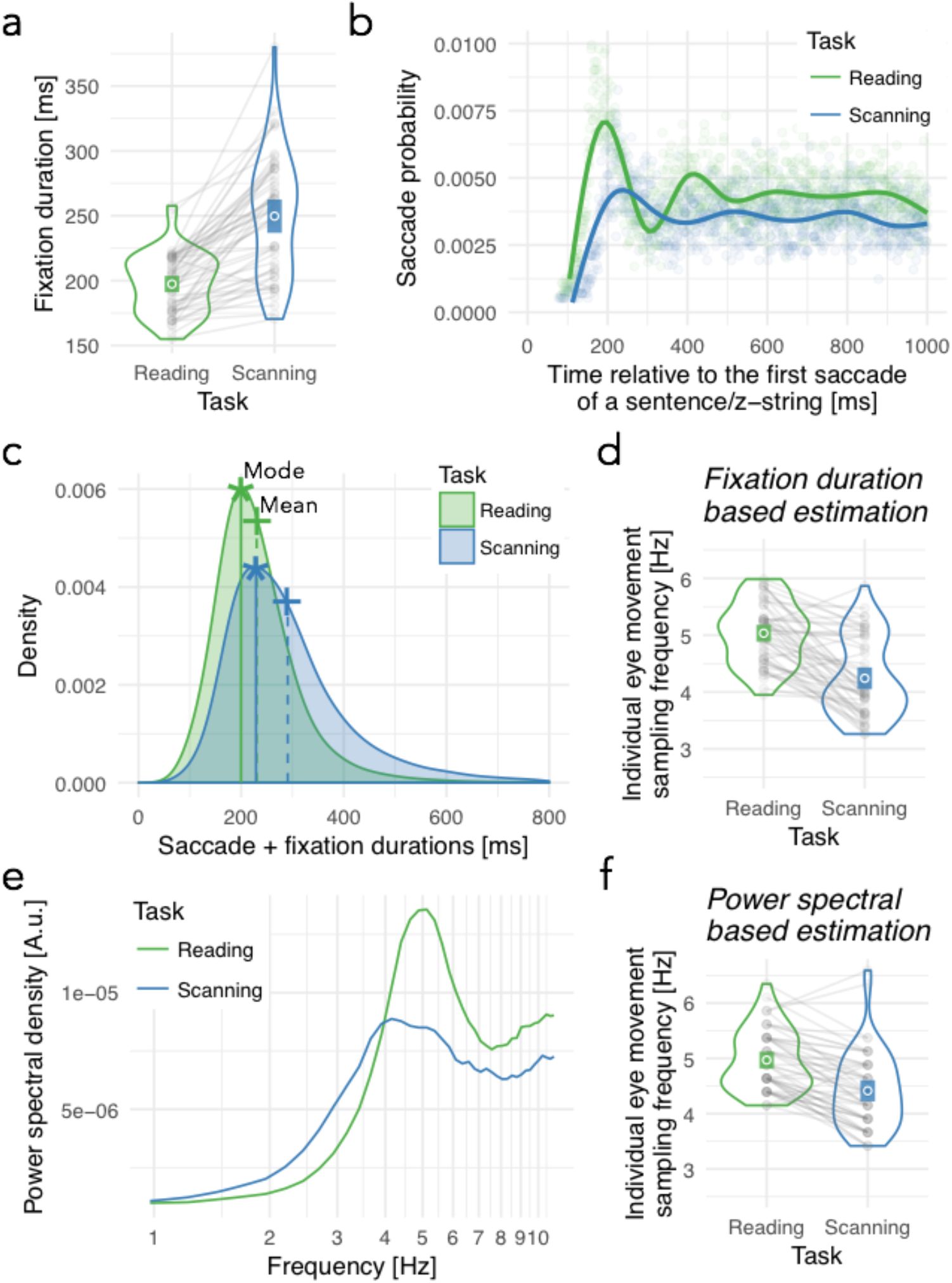
Reading-related sampling rates. (a) Subject-specific mean fixation durations from 50 participants (dots), the overall mean (circle), and confidence intervals (colored bars) while reading sentences on the Potsdam sentence corpus^10^ and scanning z-strings. Lines connect reading with z-string scanning data, per subject, to visualize effects at the single-subject level. Violin plot shows the distribution of individual means (Blue: Scanning; Green: Reading; similar in d and f). (b) Mean saccade probability (across all participants and stimuli, separated by task) relative to the first saccade of the sentence, with a non-linear regression line. (c) The sampling period t of one event was defined as the duration of a fixation plus its preceding saccade. Displayed is the distribution of these sampling periods for sentence reading (green) and z-string scanning (blue), with estimated means (‘+’ symbol and dashed lines) and modes (asterisk and solid lines). (d) Subject-specific mean sampling frequencies f (i.e., equals to 1/t) and the overall mean (crossed circle) based on the sampling periods shown in c. (e) Power-spectrum for reading and z-string scanning, estimated across all participants using Fourier transform analysis. (f) Individual peak frequencies estimated from individual power spectra and their mean (crossed circle). See Methods for details.

As a first characterization of rhythmic eye-movement patterns during reading, we plotted for each sample point after stimulus onset the probability that a saccade occurs (Fig. 1b). This analysis demonstrates distinct peaks visible at regular intervals, providing evidence that eye-movements follow a rhythmic structure in both reading and scanning. Importantly, this rhythmic pattern is more pronounced and faster during reading. Dominant sampling rates were estimated directly from fixation durations, as well as using classical frequency analysis. While the former approach is important because fixation durations are also the basis for the subsequent meta-analysis, the latter approach allows us to evaluate the validity of fixation-duration-based frequency estimation.

To estimate sampling rates from fixation durations, we first estimated sampling periods *t* (i.e., the time from the start of a saccade to the start of the next), by adding to each fixation duration (N=112,547) the duration of the preceding saccade. Figure 1c shows the distribution of all sampling periods across participants, separately for reading and z-string scanning. Note that due to the ex-Gaussian distribution of fixation durations typical for fixation duration data^21^, the mean (dashed line) overestimates the central tendency, whereas the mode (solid line) – by definition – is a better representation of the predominant sampling period (Fig. 1c). Next, we estimated an eye-movement sampling frequency *f* for each participant and condition, by dividing 1sec by the subject-specific mode of the sampling period in seconds. This revealed a higher average sampling rate for reading (5.0Hz) relative to the control task (4.2Hz; Figure 1d). This difference was significant (Cohen’s d=-1.16; *t*(49)=-8.2; p<.001) and 45 of 50 participants showed a numeric reduction of sampling frequency from reading to scanning (grey lines in Fig. 1d). We find virtually the same pattern of effects when regressive saccades are removed (i.e., when analyzing only single fixation cases; Cohen’s d=-1.0; t(49)= -6.9; p<.001; absolute values: 4.9Hz and 4.2Hz for reading and scanning, respectively) and only slightly higher values when restricting analyses to inter-word re-fixations (i.e., fixations after regressive saccades; 5.2Hz vs. 4.6Hz, respectively; Cohen’s d=-0.8; t(48)=-5.5; p<.001; note that one participant was excluded due to the absence of regressive saccades in the scanning task). Sampling rates of reading and scanning, thus, are highly similar between forward-oriented and regressive eye movements (*r*=0.6; *t*(96)=6.7; *p*<.001). Therefore, all further analyses will not differentiate between these cases. Note that estimating sampling rates from the mean (rather than mode) of fixation durations results in lower rates for reading (4.5Hz) and scanning (3.7Hz). This results from an overestimation of the central tendency by the mean in right skewed distributions (see Fig. 1c) and indicates that this procedure would be inadequate.

Finally, power spectra of reading vs. z-string scanning were estimated using canonical frequency analysis. For each task, we created a time series (resolution 1,000Hz) starting with the first saccade of the first participant and ending with the last fixation of the last participant, with a ‘1’ at the exact time of saccade onset and ‘0’ elsewhere. Note that saccade onsets are the appropriate event for generating this time series, as they are the re-occurring event and can be measured with high accuracy^22^. Subsequently, power spectra of these task-specific event timecourses were estimated via Fourier Transform to visualize periodic signal components across subjects (see Methods). Corroborating the results of the first analysis approach, a prominent peak was found at 5Hz for reading and a somewhat less pronounced peak at ∼4Hz for scanning (Figure 1e). To compare these estimates between reading and scanning, we next estimated separate power spectra for each participant. Individual peaks were retrieved, averaged (Figure 1f), and statistically compared. This analysis reproduces the sampling frequencies estimated from the mode of fixation durations, with frequencies of 5.0Hz and 4.4Hz for reading and scanning, respectively (Cohen’s d=-1.12; *t*(49)=-7.9; p<.001). There was a high correlation between the two analysis approaches (reading: *r*=.80; *t*(48)=9.3; *p*<.001; scanning: *r*=.62; *t*(48)=5.5; *p*<.001), which underscores the validity of sampling-duration-based frequency estimations.

To summarize, a quantitative frequency-domain characterization of eye-tracking data shows that the predominant sampling frequency during reading in German, across participants, is found at ∼5Hz. This frequency representation of the reading process falls squarely within the boundaries of the predominant modulation frequencies of 4.3-5.5Hz determined for speech signals across languages^5,6^ which in turn have a clear reflection in the neuronal response to speech^3^. We observed the ∼5Hz peak during reading using two different analysis strategies, i.e., when estimating sampling frequencies from saccade and fixation durations and when analyzing the sequence of saccade events in the frequency domain. Attentive scanning of z-strings shows highly similar scan path characteristics compared to reading^18,19^, but a significantly lower sampling frequency at ∼4.2Hz, convergent with findings from non-linguistic attentional reorienting tasks^15,23^.

An analysis of the pupil response in this same dataset had previously indicated higher cognitive effort during reading as compared to z-string scanning^17^. This finding most likely reflects the additional involvement of reading-specific and linguistic processes, like lexical-semantic access, beyond the oculomotor sampling itself. Thus, the specific sampling rate observed for reading is unlikely to be driven exclusively by (perceptual or cognitive) features of the stimulus. In light of the overlap with the rate of spoken language, we tentatively propose that the observed sampling rate of ∼5Hz may reflect functional constraints imposed by the interface nature that the process of reading has between visual and linguistic processing (which developed primarily based on spoken language). We speculate that the brain’s language systems impose the cortical rate at which speech is produced and perceived onto oculomotor programming systems exclusively during reading, possibly to optimize language-related information processing.

This hypothesis predicts that the overlap of reading and speech rates should generalize across languages and writing systems. On the other hand, writing systems differ substantially between languages^11,24^, and even within writing systems, the mapping from orthography to meaning differs between languages^25^. For example, the letter *a* in *cat* vs. *ball* maps onto two different speech sounds in English, whereas it maps onto the same sound in the German translations of these words (*Katze* vs. *Ball*). This letter-to-sound correspondence strongly influences reading acquisition^26^, so that among the alphabetic writing systems, opaque orthographies (writing systems like English with inconsistent letter-to-sound correspondences) are associated with lower reading accuracy during the first years of learning to read. These differences would be suggestive of cross-linguistic differences in the frequency at which written text can be sampled, and recent experimental evidence like the observation of longer fixation durations for Chinese as compared to Finnish or English^11^ seems to support this prediction.

## Results: Cross-linguistic meta-analysis of reading rates

To investigate the language generality of the alignment between speech and reading rates, we conducted a meta-analysis of sampling frequencies during reading in 14 different languages, based on 1,420 fixation duration estimates extracted from 124 original studies published between 2006 and 2016 (see *Methods* for selection criteria). In addition to this cross-linguistic comparison, we examined (a) possible differences between character-based vs. alphabetic writing systems and (b) the effect of letter-to-sound correspondence among alphabetic writing systems. Also, we explore (c) the cross-linguistic correlation between eye-movement sampling frequencies and language-specific peaks of the speech modulation spectra, and (d) the association between reading rates and information density (linguistic information per syllable^27^) across languages.

All studies selected for inclusion reported mean fixation durations. However, as discussed above, mean fixation durations are not a valid representation of the predominant sampling duration in fixation data and accordingly not the preferred basis for calculating the sampling rate of reading. We used 29 full empirical datasets to develop a transformation function that allowed us to estimate the mode from the mean fixation durations reported in the original publications. In brief, this involved fitting ex-Gaussian distributions to the empirical distributions of these datasets, retrieving distributional parameters (including mean and mode), and on this basis optimizing a regression-based transformation that estimates the mode from the mean (see *Methods* and *Supplementary Methods* for details). For the meta-analysis, mean durations were extracted from published studies and transformed to the mode. The sampling period *t* (interval from saccade onset to end of following fixation; see above) was obtained by adding an estimate (the mode saccade duration from Study 1; 29ms) to the mean fixation duration. Lastly, the sampling frequency was calculated as *f*=1/*t*.

### Fixation durations and sampling frequency: Descriptive statistics

Figure 2 shows that the majority of mean fixation durations derived from the reading studies were between 200-300ms (upper panel), which transforms to mean sampling frequencies between 3.9-5.2Hz (lower panel). Note that languages with only one original study (Arabic, Italian, Polish) were excluded. As expected, the majority of studies was conducted in English^28^. 10 of the 14 languages in our meta-analysis fall between the minimum (4.3Hz) and the maximum (5.5Hz) of previously reported^5,6^ language-specific peaks of the speech amplitude modulation spectra (see Figure 2, lower panel, dashed lines). The remaining four languages fell within the range of one standard deviation around the language-specific speech peaks (Figure 2, lower panel, dotted lines). Considering the language-specific confidence intervals, only for Chinese can we be confident that the sampling rate is lower than the range of the speech amplitude modulation spectra^5,6^. Of the 1,420 individual sampling rate values derived from the studies included in our meta-analysis, only 3.0% fell below, and only 0.3% were above the range of one standard deviation of the mean (see violin plot in the lower panel of Figure 2). Nevertheless, the mean sampling rate of reading observed when averaging across all languages is at the lower bound of the speech modulation range, i.e., at 4.3Hz (Figure 2, lower panel).

**Figure 2.**
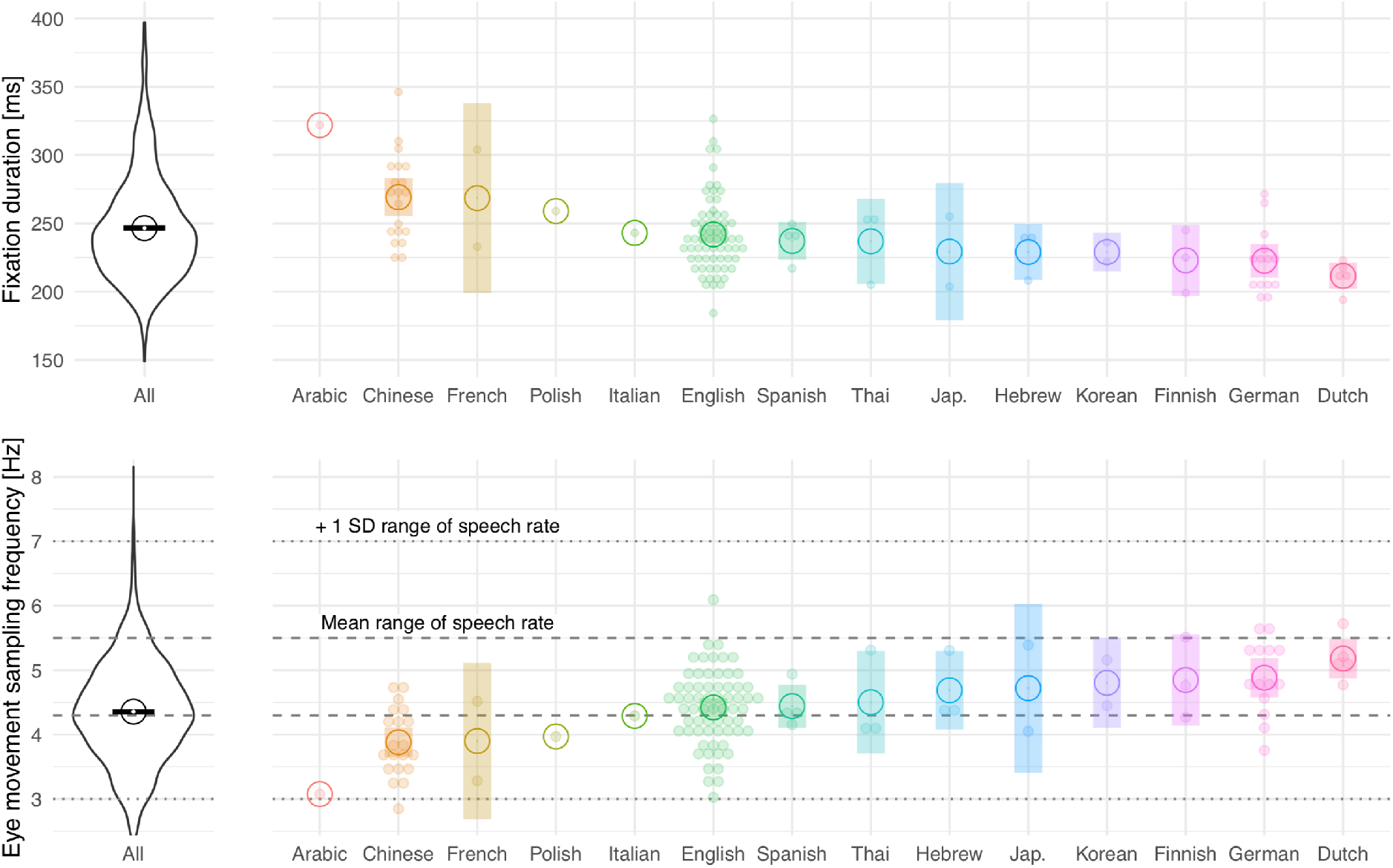
Meta-analysis of reading-related sampling rates. Fixation durations (upper panel) and corresponding eye movement sampling frequencies (lower panel) for 14 different languages. Violin plots (left) represent the respective distributions of all 1,420 duration/frequency values extracted from the included studies, independent of language. Bars reflect confidence intervals, and circles reflect the mean. In the right panel, each dot reflects one study (mean number of fixation durations per study: 12.4); Bars reflect confidence intervals, and circles reflect the mean across studies for each language. In the lower panel, the dashed lines represent the range of the means of the peak amplitude modulation spectrum that was empirically determined for speech in different languages in independent work^5,6^. The dotted lines represent the range between the lowest mean minus one standard deviation and the highest mean plus one standard deviation for the same data (that was manually read out from Figure 3c in Ref.^5^ and from Figure 7 in Ref.^6^). For Arabic, 1 study/ 12 fixation durations are available, Chinese 20/205, Dutch 5/45, English 65/965, Finnish 3/21, French 2/3, German 14/48, Hebrew 3/28, Italian 1/1, Jap. 2/12, Korean 2/39, Polish 1/1, Spanish 4/10 and Thai 3/30.

### Effect of writing system on sampling frequency

The observed cross-linguistic differences, arguably, are related to different language characteristics. One plausible hypothesis is that the higher perceptual complexity of character-based scripts (as opposed to alphabetic scripts^24^) may modulate the rate at which written text is sampled. Figure 3a shows that the eye-movement sampling frequency is significantly lower for Chinese (the only character-based language included; n=205 estimated sampling rates from 20 studies; mean: 3.9Hz) than for alphabetic languages (n=1,215 sampling rates; 97 studies; mean: 4.5Hz; effect size estimate (Est) of difference: -.70Hz; Standard error (SE): .13; *t*=5.2; see *Methods* for details on linear mixed-effects modeling).

**Figure 3.**
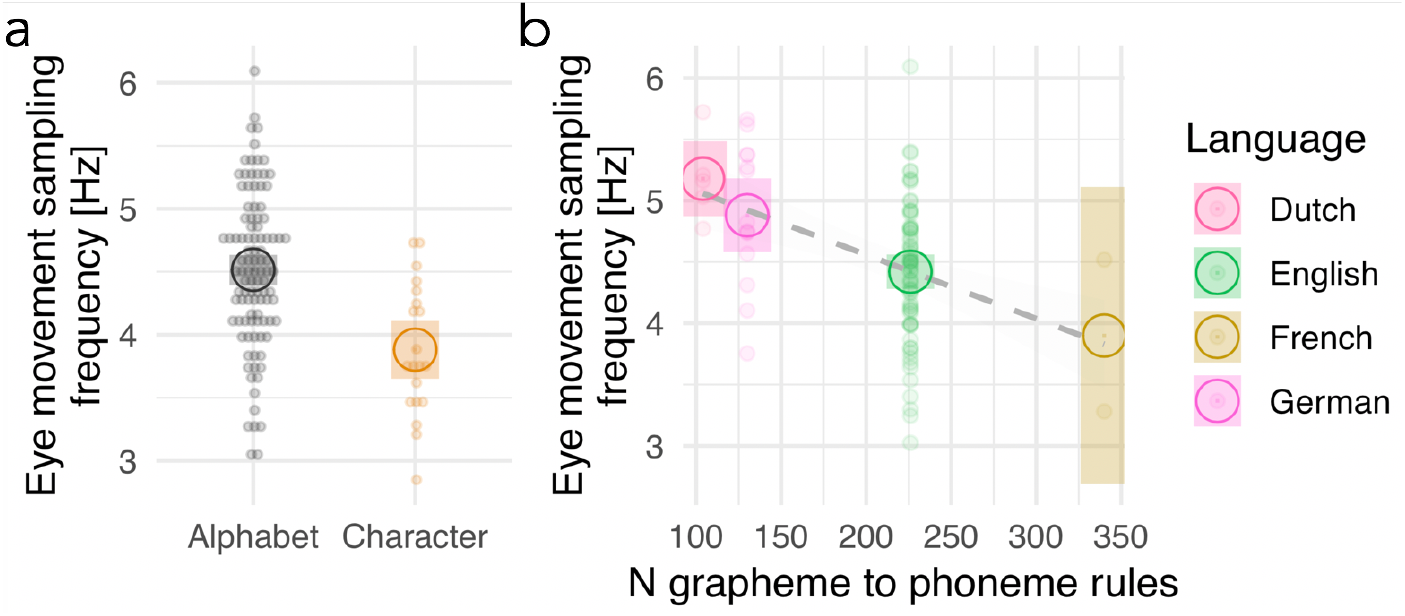
Comparison of writing systems. (a) Character vs. alphabetic contrast, including 205 fixation durations from 20 Chinese reading studies (brown) and 1,215 fixation durations from 97 studies of reading in alphabetic languages (grey). (b) The effect of language transparency/opacity. Only studies from alphabetic languages for which the number of grapheme-to-phoneme rules could be quantitatively estimated from published computational models (see *Methods*) were used (four languages with a total of N = 1,025 fixation durations). Dots reflect each study, and crossed circles reflect the mean across studies for each language. The dashed line in (b) represents the approximation of the language transparency/opacity effect based on a linear regression.

### Effect of orthographic complexity on sampling frequency

Within alphabetic languages, one plausible hypothesis is that the orthographic difficulty of writing systems influences the speed of sampling the visual input^29^. To examine this, we quantified orthographic difficulty as a continuous predictor representing the number of grapheme-to-phoneme conversion rules^30^ as defined by computationally implemented dual-route models of visual word recognition^31^. Graphemes are letters or letter combinations that map onto one or multiple speech sounds (phonemes); for illustration, remember the above example of mapping the grapheme *a* onto one (Katze/Ball) vs. two (*cat*/*ball) phonemes*, requiring one vs. two rules. To date, computational implementations are available for only five out of the nine alphabetic languages included in this meta-analysis, which restricts this test to English, French, German, and Dutch (with n = 965, 3, 48, 45 data points, respectively; Italian, the fifth language, was excluded due to lack of sufficient data points). A detailed comparison of these language-specific model implementations can be found in Ref.^25^. Figure 3b demonstrates that less transparent writing systems (operationalized as more grapheme-to-phoneme rules) elicit significantly lower sampling frequencies (Est: -.10Hz; SE=.03; *t*=3.0). Highly transparent orthographies (German, Dutch) produce relatively fast sampling rates around 5Hz (Fig. 3b).

### Effects of speech rate and information density on sampling rates

Lastly, we explored the effect of cross-linguistic differences in speech rate^5,6^ and information density (information per syllable^27^) on the observed eye-movement sampling rates (see *Methods*). To control for the strong effects of orthographic differences on sampling rates (as above), linear mixed models were calculated that also included the factor alphabetic vs. character-based script (effect size estimates/Est in both models<-0.57Hz; SE<0.15; *t*>4). Neither the between-language differences in speech frequencies (Est: -0.03; SE=.03; *t*=0.8) nor information density (Est: - 0.03; SE=0.05; *t*=0.7) showed significant effects on the eye-movement sampling rate (all analyses including Chinese, Dutch, English, French, Japanese, and Spanish).

As further exploration, we investigated the relationship between speech and reading rates within alphabetic languages for which estimates of orthographic complexity (grapheme-to-phoneme rules) could be taken into account (English, French, Dutch). This analysis also failed to produce significant effects (Est.:0.06; SE=0.06; t=1.0). Still, the result indicated a positive relationship between peak speech modulation rate and eye-movement sampling rate. Note that we report this analysis despite its low statistical power (with only three languages), to motivate future investigations of the relationship between speech and reading rates.

The meta-analysis (i) replicates the results obtained for German in the first section, (ii) shows that eye-movement sampling frequencies of most languages fall into the range of the peaks of speech amplitude modulation spectra (4.3-5.5Hz)^5,6^ determined in independent research, and (iii) shows a systematic modulation of reading rates by the perceptual difficulty of orthographic systems. We found similar average sampling rates for languages of comparable orthographic transparency levels (e.g., German and Dutch) and highest reading rates (∼5Hz) in transparent (i.e., relatively easy-to-process) writing systems. Our tentative proposal that the linguistic processing systems underlying speech production and comprehension provide the temporal frame that ‘drives’ the oculomotor machinery during reading would predict a direct relationship between the rate of speech production and the sampling rate of reading. In the present meta-analysis, this proposal could only be tested cross-linguistically and using highly aggregated data, and we found no robust support for this proposal. However, these analyses included small sample sizes as they were limited by the number of languages included. To investigate this proposal in more detail, we next conducted two new studies that examine the existence of associations between speech and reading rates at a subject-by-subject level.

## Results: Association of individual differences in speech and reading rates

We tested the correlation between peaks in the speech modulation spectra of individual speakers and their eye-movement sampling rates during reading in two experiments. First, we tested 48 learners of German (Study 3), as we expected to observe higher variabilities in both measures in non-native language learners than in native speakers^32^ and a more direct relationship between speaking and reading (similar to letter-by-letter reading in beginning readers^33^). We recorded eye movements from each participant while reading German sentences (implemented analogously to the reading task in Study 1) and a speech sample based on a ‘small talk interview’ (22 questions, on average 18 minutes of speech per participant, range: 6 to 28min). Also, we controlled for individual differences in reading proficiency statistically by adding a standard measure of reading skill^34,35^ (see *Methods*). The eye-movement sampling frequency was estimated based on the fixation durations (i.e., as in Experiment 1), and the speech modulation spectrum was examined analogous to previous reports^6^.

Figure 4a shows the average speech modulation spectrum across participants (black line), with a peak at 4.2Hz, and individual spectra from all participants (gray lines). As expected for language learners, the peak of the spectrum was below that of native speakers (compare Figure 1) and on the lower border of the cross-linguistic range of mean speech rates^5,6^ (compare Figure 2). Nevertheless, all participants produced peaks between 3Hz and 6Hz (Figure 4b), i.e., within the previously reported^5,6^ range of the standard deviations around the language-specific mean peaks (dotted line in Figure 4c). The peaks of individual speech modulation spectra and eye movement sampling frequencies were in a comparable range (Figure 4b; confirmed by a significant equivalence test^36^: *t*(47)=-2.0, *p*=.03) and positively correlated (Figure 4c; Effect estimate/Est=0.32; SE=0.15; *t*=2.1; *p*=0.04). Note that this correlation effect was estimated while controlling for individual differences in reading proficiency (Figure 4d; Est=0.032; SE=0.016; *t*=2.1; *p*=0.04) by calculating a linear model that estimates the individual eye-movement sampling rate with speech modulation rate as predictor.

**Figure 4.**
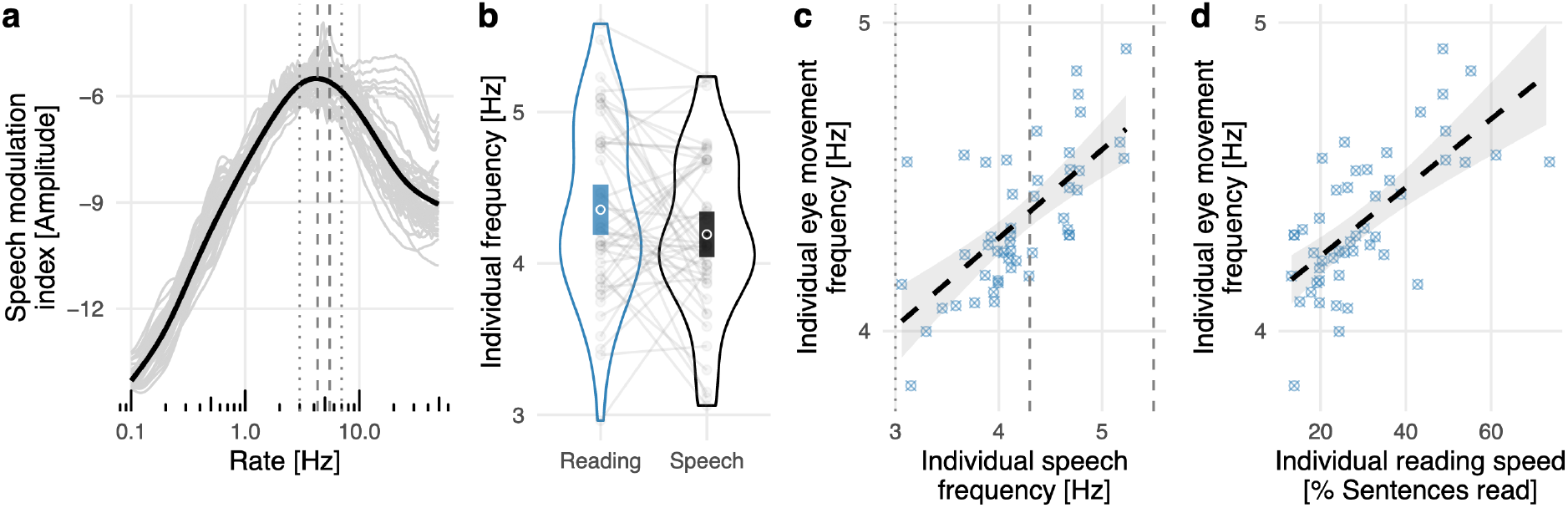
Relationship of speech and reading rates in non-native German speakers. (a) Speech modulation spectrum from 48 non-native speakers of German. Y-axis: speech modulation index^6^; X-axis: speech modulation rate. For additional comparison, we present the mean range (dashed lines) and standard deviations (dotted lines) of the speech amplitude modulation spectra across languages, which were read out from Figure 3c in Ref. ^5^ and from Figure 7 in Ref. ^6^. (b) Eye movement sampling frequency in reading and the mean amplitude modulation spectrum in speech, for each participant. Lines connect the reading and speech frequencies of each individual, the violin plots represent the distribution of the data, bars represent the standard error of the mean and circles reflect the mean. (c) Positive correlation of the individual peaks of the speech modulation spectrum (x-axis), reflecting each participant’s speech rate, with the eye-movement frequency (y-axis) from the same participants. (d) Correlation between the eye-movement frequency (y-axis) and a paper-pencil based reading score (x-axis) reflecting a positive association of the eye-movement sampling rate and reading performance. Note that in (c) and (d) we present the individual sampling frequencies corrected for either reading skill and speech frequency, respectively, based on predictions from the fitted linear regression models used for statistical analysis.

In a second, pre-registered study (Study 4), we assessed the relationship between speech modulation spectrum peaks and eye movement sampling frequencies in a group of 86 native speakers of German (see Method; preregistration: https://osf.io/mjhkz). We replicated the finding that the peaks of the speech modulation spectra and eye movement sampling frequencies were in the same range (Figure 5a; equivalence tests for left and right eye: t’s>3.9, p’s<.001). The pre-registered correlation analysis showed a small positive, albeit not significant relationship between eye-movement sampling and speech modulation rates (Est.=0.07; SE=0.04; t=1.8; p=0.08). For further exploration (i.e., non-registered post-hoc analysis), we separately investigated and compared four subgroups created by a 2×2 combination of reading speed and reading accuracy. Specifically, we implemented a median split based on reading speed measured with a standardized German reading test (adult version of the SLS^35^; fast vs. slow: Median: 78% vs. 60%) and, orthogonal to this, divided the sample based on their sentence comprehension accuracies in the eye-tracking experiment (errors present vs. absent: Median 0% vs. 15%). Only readers with the lowest skill level, i.e., slow and low comprehension performance, showed a robust positive association (N=21; Figure 5b, bottom right; Est.=0.33; SE=0.10; *t*=3.4; *p*=0.002). None of the other groups showed a significant correlation, resulting in a reliable interaction effect (Est.=-0.05; SE=0.02; *t*=2.8; *p*=0.005) which was also present when the percentage of regressions, skipping, and single fixation probabilities (see Table 1) were added as covariates to the model. Note that the low-reading skill group had a lower reading speed than the ‘slow only’ group that produced no errors (Est.=0.08; SE=0.04; *t*=2.0; *p*=0.049), but still had a substantially higher reading speed compared to the non-native readers from Study 3 (Est.=0.57; SE=0.04; *t*=14.42; *p*<.001; for general eye-movement characteristics of both experiments see Table 1). In sum, we replicated the results of Study 3 selectively by demonstrating that the correlation between reading and speaking rates is limited to (here native German) speakers with low reading skills.

**Figure 5.**
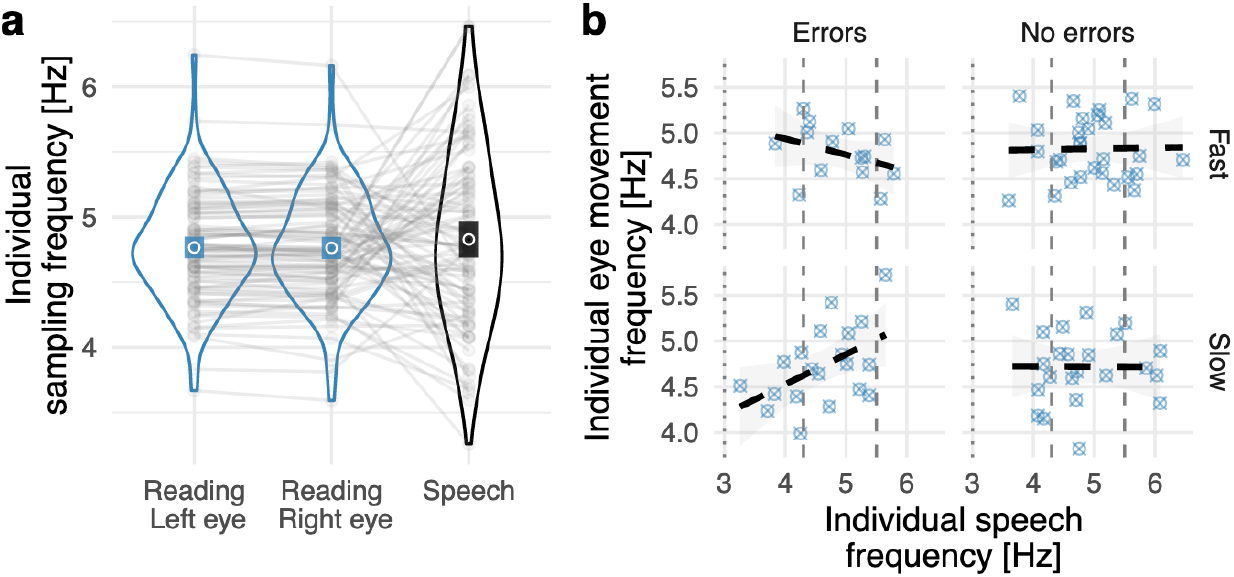
Relationship of speech and reading rates in 86 native German speakers. (a) Eye movement sampling frequency measured during reading for the left and right eyes (left and center columns) and the mean amplitude modulation spectrum of samples of spoken speech (right column). Grey dots represent individual data points from all participants; lines connect the reading and speech frequencies of each participant. The violin plots represent the distribution of the data, filled bars represent the standard error of the mean, and circles reflect the mean. (b) The correlation between the speech modulation spectrum (x-axis) with the eye-movement sampling frequency (y-axis). The four panels represent performance subgroups depending on reading speed (slow vs. fast; median split) and whether they produced errors in an independent standardized reading test (see Methods for details).

**Table 1.** Reading speed, reading comprehension, and basic eye tracking measures (fixation durations, skipping probability, single fixation cases, and percentage of regressions) for Study 3 and Study 4. For Study 4, data are presented separately for the four performance-based subgroups. All values reflect means and standard deviations.

|  | Study 3 | S. 4 Fast; No errors | S. 4 Fast; Errors | S. 4 Slow; No errors | S. 4 Slow; Errors |
| --- | --- | --- | --- | --- | --- |
| Reading speed (SLS test; % sentences read) | 26.0 (10.4) | 79.0 (6.5) | 78.9 (5.3) | 58.0 (11.1) | 53.6 (9.0) |
| Reading speed (experiment; in sec.) | 4.8 (2.3) | 2.4 (0.7) | 2.2 (0.8) | 2.4 (0.5) | 2.7 (0.6) |
| Reading comprehension (experiment; % errors) | 20 (40) | 0 (0) | 17 (4.4) | 0 (0) | 19 (14.5) |
| Fixation duration (right eye, in ms.) | 223 (29) | 192 (20) | 188 (15) | 196 (21) | 198 (20) |
| Fixation duration (left eye, in ms.) | 225 (30) | 192 (20) | 189 (15) | 197 (21) | 199 (20) |
| Skipping (%) | 7.5 (5.8) | 19.8 (7.9) | 24.0 (10.8) | 15.7 (6.9) | 14.7 (6.6) |
| Single fixation cases (%) | 35.5 (14.6) | 52.3 (9.1) | 54.1 (10.2) | 56.6 (9.2) | 52.7 (9.4) |
| Regressions (%) | 10.9 (4.5) | 12.1 (5.3) | 9.3 (6.9) | 8.6 (4.9) | 10.6 (6.0) |

*Note.* Statistical comparisons of the four groups from study 4 (2x2, error by speed groups interaction linear regressions models) yielded a significant main effect of reading speed group (slow vs. fast) for SLS-based reading speed (Estimate: 0.53; SE = 0.22; $t(84) = 2.4$ , $p = 0.017$ ; all other effects $t$ 's < 1.7) and an obvious group difference on the percentage of errors (0 vs. ~18% errors). For eye-tracking measures, the only significant effect was a main effect of reading speed group on the percentage of skipping (Estimate: -0.09; SE = 0.03; $t(84) = 3.4$ , $p < 0.001$ ); no other main effect or interaction effect reached significance (all $t$ 's < 1.94).

## Discussion

This (to our knowledge first) frequency-based investigation of eye-movements during reading shows that reading operates in a generally comparable frequency domain as the production and perception of natural speech. We first reproduced in a frequency-domain analysis previous insights based on fixation duration measures^17–19^, i.e., that eye-movements sample text with a higher rate than during comparable, cognitively less challenging non-linguistic tasks. More importantly, we demonstrate that the sampling frequency of reading lies within the range of previously observed speech rates for one language, German. Next, by integrating across languages data from 124 empirical studies, we show that eye-movement sampling varies between ∼3.9Hz and ∼5.2Hz, indicating a higher variability than previously assumed. While it was generally believed average fixation durations are similar even for very distinct orthographies like Chinese and English (Rayner^12^, p.1461), our meta-analytic results that show significantly higher sampling rates for alphabetic compared to character-based writing systems. However, average speech rates have been shown to vary more narrowly around 5Hz across languages (i.e., Ref.^5^: 4.3-5.4Hz; Ref.^6^: 4.3-5.5Hz), a range that would exclude the lower frequencies we observed for reading. Our meta-analytic findings indicate that this might result from differences in the complexity of the underlying orthographies (e.g., character vs. alphabetic), such that more computationally ‘difficult’ orthographies might slow down reading relative to highly transparent alphabetic orthographies.

Subsequently, we demonstrate in two independent empirical studies that second language learners (of German) read in a lower frequency range than native readers (∼4.3Hz vs. ∼4.7Hz) and that only language learners and low skilled native readers show a positive correlation between individual reading and speech rates. Combined, these results suggest that reading, i.e., an internally controlled visual-perceptual process involving sophisticated oculomotor programming, is remarkably well temporally aligned with the rate at which spoken language is produced (and perceived). We tentatively suggest that this observed association between speech and reading supports the existence of fundamental perceptual principles underlying the temporal structure of linguistic information processing, irrespective of modality^16^.

Text is a temporally stable visual stimulus. However, our eye-movements impose temporal structure onto the linguistic input when sequentially sampling a text. The reading process - including the oculomotor programs - thus serves as an interface between a stable external percept and linguistic processing systems optimized for analyzing sequential speech input. The observation of faster sampling rates during reading as compared to parsing non-linguistic letter strings (Study 1) indicates that sampling rates are not exclusively driven by the physical layout of the stimulus or by the cognitive effort of processing the stimulus (in which case they should have been slower). We tentatively propose that neural processors dedicated to the linguistic analysis of speech impose their preferred timing onto the process of reading. Evidence for the principled possibility of such internally driven entrainment of reading comes from the observation that manipulating the speed of ‘inner speech’ during reading has a causal effect on reading speed^37–39^.

Our meta-analysis demonstrates that the sampling rate of reading varies between languages – but falls within the range of speech rates identified in cross-linguistic studies^5,6^. The meta-analysis also shows higher sampling rates for transparent vs. opaque orthographies, which converges with transparency effects within languages^29,40^ and cross-linguistic studies investigating reading development^26,41,42^. Direct associations between reading and speech rates could not be established in the meta-analysis given the small number of languages for which all necessary parameters were available. Empirical Studies 3 and 4 show this relationship on a subject-by-subject level, however only in less-skilled readers. This suggests that increasing reading expertise makes the tight control of reading by linguistic processors in the brain obsolete.

The differential coupling of speech and reading rates in low-skilled but not high-skilled readers may also result from other phenomena well-established in reading research, like word skipping, para-foveal preprocessing, and re-fixations. Reading is not merely a sequence of word-to-word fixations. From time-to-time, we skip words as a result of parafoveal pre-processing^43,44^, which describes visual word recognition based on low acuity visual information from parafoveal regions of the retina. Also, words are sometimes fixated multiple times, e.g., to correct perceptual errors after suboptimal landing at the beginning or end of a word^14,18,45^ or when semantic inconsistencies must be resolved by re-reading^46,47^. We^14^ and others^10^ have shown that overall probabilities for word skipping and multiple fixations on a word are comparable (∼20%) when reading the sentence materials used here. However, low-skilled readers show lower skipping rates (reflecting reduced parafoveal preprocessing^48–50^) and more re-fixations on the same word^13^. Thus, readers with lower skills focus more on fixated words and their components (letters, syllables)^51^, indicating a greater alignment between the phonological properties of the words and the eye-movements they elicit during reading^33^, suggesting that low-skilled readers sample written text with a temporal resolution close to the speech processing rate. In contrast, the faster reading rates of fluent readers indicate that they utilize the static nature of text better by processing the fixated word and following words (based on parafoveal vision) within one ‘sample’.

The specific characteristics of fixation behavior and text presentation during reading can also provide context for another intriguing phenomenon, i.e., the significantly lower sampling rates in character-based than alphabetic writing systems (while overall reading times for sentences with the same content are comparable between the two writing systems^11^). Here, fewer fixations per sentence are needed to sample the entire stimulus, while the increased perceptual complexity and information density^11^ lead to longer fixation durations, relative to alphabetic languages. In the non-linguistic control task, participants were presented with stimuli consisting of many repetitions of the same letter. In this case, information density and perceptual complexity are low compared to all real-language stimuli. Nevertheless, we observed longer fixation durations in the z-string scanning task, which may indicate the presence of qualitatively different cognitive processes compared to reading.

The frequency representation of reading-related eye-tracking data that we advance here can be construed as ‘nothing but’ a transformation of fixation and saccade duration data. This transformation also comes at the cost of zooming out to a ‘meso-level’ representation of the data^1^, at which we rely on aggregated data (i.e., only one data point per participant), which is against the trend in eye-movement research of focusing on investigating single words and using regression methods for detailed analysis of, e.g., the influence of word characteristics^52^. Still, the frequency perspective proposed here provides a novel perspective onto component processes of reading as an interface between linguistic and orthographic processing. This new approach to reading research opens up several interesting new research questions. For example, it becomes possible to compare reading behavior more directly to evidence from other measurement modalities, such as oscillatory brain activation data^53,54^, and to other cognitive-psychological domains, such as attention^15,55^, which typically do not have the advantage of exact duration measurements for different events of interest (e.g., during covert attention). Maybe most importantly, the frequency perspective on reading offers direct links to several neurodynamic phenomena in speech perception^5,6^, including the observation that dyslexic children^56,57^ and adults^58^ show altered cortical tracking of speech signals in the oscillatory domain.

In conclusion, we show that during reading, our eyes ‘sample’ written text in the same frequency range in which speech is produced and perceived, which suggests that extracting information from linguistic stimuli follows a similar temporal structure in time irrespective of modality. A plausible mechanism is to assume that linguistic processing has a preferred cortical rate of information uptake and thus acts as an internal temporal driver for eye-movements elicited during reading. Thus, eye-movements in reading are utilized as a temporal interface between a stable physical stimulus – written text – and brain systems that have evolved to process signals whose temporal structure is constrained by the characteristics of our vocal tract^1^. However, our empirical data also demonstrate that a direct coupling between speech and reading rates is only present in persons with low reading skills, which calls for future work to clarify the mediating role of reading expertise for the temporal relationship between speech processing and reading rates. We suggest that the novel frequency perspective on reading adopted here opens up new research paths, such as understanding slow or impaired reading, second language learning, or more directly investigating the commonalities and differences between reading and other cognitive processes.

## Methods

### Study 1, 3 and 4, Participants

Fifty (13 male; 18–47years old; M=24years; students at University of Salzburg) native speakers of German participated in Study 1, forty-nine (13 male; 18–74years old; M=24 years) non-native German speakers participated in Study 3, and eighty-six (36male; 18–53years old; M=25years; five had to be excluded based on preregistered outlier correction boundaries for both speech and reading rates; +-3 standard deviations) German speakers participated in Study 4 after giving informed consent according to procedures approved by the respective local ethics committee. For Study 1, see our original publication of this dataset^17^ for more details. Note that relative to the previously published report, one participant was added. For Study 3, participants with varying mother tongues (Arabic, Azerbaijani, Bulgarian, Chinese, English, Farsi, French, Georgian, Indonesian, Italian, Japanese, Persian, Russian, Serbo-Croatian, Spanish, Turkish, Ukrainian, Hungarian, Urdu, and Uzbek) and, for Study 4, native German participants were recruited on the campus of Goethe University Frankfurt as part of a larger study. Also note that six participants from Study 3 became literate without the acquisition of an alphabetic script. The power estimation for the fourth and final study resulted at sample size of 90 while assuming the effect size from study 3 and a power of .9 (see preregistration for more details: https://osf.io/mjhkz).

### Procedure Study 1

Movements of the right eye were tracked with a sampling rate of 1,000Hz (Eyelink 1000, tower mount system; SR-Research, Ontario, Canada). We used a forehead and chin rest to fixate the head of participants at a distance of 60cm from a 21” CRT screen. In the reading task, we used the Potsdam Sentence Corpus^10^ (PSC) which consists of 144 sentences and a total of 1,138 words. Participants were instructed to read silently for comprehension, which was controlled by simple comprehension questions after 38 of the 144 sentences.

As a non-linguistic control task, participants performed a z-string scanning task using stimuli in which all letters of the sentence corpus were replaced by the letter z (preserving letter case, punctuation, and word boundaries. Participants were instructed to visually scan the meaningless z-strings as if they were reading; for obvious reasons, no comprehension questions were administered in this condition. Z-string scanning has been used as control task in previous studies^17–20^. While it is difficult to find a reasonable control task for reading (see, e.g., Discussion in Ref.^18^), z-string scanning proved to be interesting because participants produce similar scan path patterns (i.e., similar number of fixations) as when reading^17–19^.

Interestingly, while z-string scanning produced longer mean fixation durations than reading, the pupil response indicated higher cognitive effort in reading, in the dataset used here^17^. We take this dissociation between cognitive effort and reading time as evidence for the operation of reading-specific cognitive processes that go beyond mere attentional processes.

In each task, a 9-point standard calibration was performed before the 10 practice trials, before the experimental trials, and after a break halfway through the experiment. A calibration was considered accurate when the mean error was below .5° of visual angle. Visual stimuli were presented in black letters (mono-spaced, bold Courier New font; 14 pt., width ∼.3°) on white background with a 1,024 × 768 pixel resolution and a refresh rate of 120 Hz, using Experiment Builder software (SR Research, Ontario, Canada). In both tasks, a trial started when an eye-fixation was found at a dot presented 100 pixels from the left margin of the monitor, at the horizontal level of the fixation cross. For this fixation check, real-time analysis of eye-tracking data was used to present the sentence only when a fixation of at least 100ms was identified on the position of the dot. If no fixation was registered on the dot for 10 seconds, a re-calibration procedure was initiated. Following the fixation check, the stimulus (i.e., sentence or z-string) appeared, with the center of the first word presented at the position of the fixation dot. As a consequence, participants always fixated the center of the first word of the sentence first. Stimulus presentation was terminated when participants fixated an X in the lower right corner of the screen after the sentence was read. As noted, in about 25% of sentences, the presentation was followed by a comprehension question to assure that participants processed sentences semantically. This procedure was practiced in ten trials prior to the main experiment.

### Procedure Study 3 and Study 4

Eye movement measurements during reading were acquired using the same stimulus materials and experimental procedures as in Study 1, with three exceptions: We used a desktop-mount eye tracker, a horizontal 3-point calibration procedure, and we did not implement the z-string scanning task. All other parameters were unchanged. For Study 4, we not only measured the right eye but also the left eye (i.e., binocular measurement). To acquire a speech sample from each participant, we conducted a brief interview in German involving 22 questions about, e.g., last weekend’s activities (see the full list of questions in the Supplementary Methods). Speech was recorded with the Audacity software (Version 2.1.3; https://www.audacityteam.org/) on a standard computer.

## Data analysis

### Fixation durations

The first word of each sentence was excluded from analyses, since the first word is known to be contaminated by stimulus onset effects. A total of 994 words were analyzed per subject. For each participant, all fixation durations from all analyzed words were extracted. Words with fixation durations shorter than 60ms and longer than 1,000ms and saccade durations longer than 80ms were removed from the analysis (3.1% of the data) since they likely reflect machine error. On the basis of the remaining fixation durations, each participant’s individual mean fixation duration was calculated, separately for the reading and scanning tasks. To account for the well-known fact that eye fixation data have an ex-Gaussian distributions (see Figure 1c) data were log-transformed resulting in a normal distribution (Kolmogorov-Smirnov test not significant; *D* < .14; *p* > .7).

### Estimation of the sampling frequency

To estimate the sampling frequency of eye movements in reading, a repetitive event needed to be identified. We defined the time between the onset of a saccade and the onset of the following saccade as the *sampling period*, which can be transformed into a frequency value. Note that we used the EyeLink eye-tracker’s built-in saccade detection algorithm, which in a recent comparative evaluation showed the best detection rates for saccade onsets compared to all other algorithms^22^. The distribution of the sampling period is ex-Gaussian, for both reading and z-string scanning (Figure 1c). Ex-Gaussian distributions are a convolution of a normal distribution and an exponential distribution reflecting the rightward skew. As Figure 1c shows, the central tendency is best represented by the mode, so that all subsequent calculations were based on the participant-specific mode of the sampling period (which we denote *t*). These subject-specific representations of the predominant sampling period were transformed to an individual eye movement sampling frequency (*f* = 1 / *t*).

### Power spectrum

We, secondly, performed a canonical frequency analysis by estimating a power spectrum for reading and scanning in Study 1. For Figure 1e, we estimated the power spectrum based on a time series starting with the first saccade of the first participant and ending with the last fixation of the final participant for each task. For Figure 1f, the time series was cut into participant-specific time series, so that individual peaks could be recovered for each participant for each task. The time series was implemented as a sparse sequence of zeros and ones (resolution: 1,000 entries per second), set to one at time points at which a saccade was initiated, and zero otherwise. Subsequently, a Fast Fourier Transform was used to estimate a power spectrum (power spectral density; *psd_welch* function from MNE-Python^59^ ; 0-100Hz, length of the FFT used = 4096 samples) for each event time course separately.

### Speech amplitude modulation spectrum in Study 3 and 4

In a first step, all non-participant audio signals were removed from the speech samples (i.e., interviewer questions and pauses before answers). To obtain the amplitude modulation spectrum we adapted the AM_FM_Spectra script^6^ (https://github.com/LeoVarnet/AM_FM_Spectra). The first adaptation divided the recording of each participant into speech segments of 10s length, resulting in a mean number of 110 segments per participants (range 35 to 167). In Study 4, we used 20s segments to increase the efficiency of the analysis (mean number of segments: 55; range 15 to 81). The second adaptation was an increase in the resolution of the amplitude modulation spectrum by decreasing the widths of the modulation filters from 3 to 10 per octave. After the speech amplitude modulation was estimated for each 10s speech segment, we retrieved the frequency at the peak of the modulation spectrum. Thereafter, we removed outliers by first eliminating unrealistic values lower than 2 and higher than 10 Hz, and then removed all values larger and smaller than two standard deviations from the mean. This procedure removed 3% of the data. Finally, we estimated the mean across all segments for each participant. Here we found that one participant, in both studies, had a mean amplitude modulation spectrum larger than three standard deviations from the mean of the sample; this participant was excluded from the analysis.

## Meta-analysis

We included empirical studies that report eye-tracking results from natural reading tasks, published between 2006 and 2016. These studies were identified by the search term *eye movement in* “*natural reading*” *or* “*sentence reading*” *or* “*text reading*“ in the *PubMed* (https://www.ncbi.nlm.nih.gov/pubmed) and *PsychInfo* (https://health.ebsco.com/products/psycinfo) databases. Additionally, 10 studies were manually identified (e.g., on the basis of reference lists in the identified papers). From the resulting sample of 124 articles we extracted 1,420 fixation durations, including mean fixation durations (all fixations on a word combined; 10% of the meta-analytic dataset), first fixation durations (duration of the first fixation on a word, which is most often reported in eye-tracking studies; 67% of the meta-analytic dataset), and single fixation durations (fixation duration in case a word was fixated only once, which is the predominant case for normal readers^10^; 23% of the meta-analytic dataset). A full list of all included studies can be found in the Supplementary Note. Note that the results of the above-reported experiment (Study 1) and its previous analysis^17^ were not included. This meta analytic dataset encompassed 14 different languages, with a range from one (Arabic, Italian, and Polish) to 65 (English) retrieved papers. Consistent with a general bias towards English in reading research^28^, 68% of fixation durations in our dataset were from English.

### Frequency estimation

In order to estimate the predominant sampling frequency, per published study, we have to take into account, once more, the ex-Gaussian distribution of fixation duration data. Following the general trend in the eye movement reading literature, most studies reported only mean fixation durations (with the exception of Ref.^21^, for which in addition the fitted ex-Gaussion paramaters were reported). For the purposes of the present meta-analysis, we developed a transformation function that allowed us to estimate the mode from the mean fixation durations reported in the original publications. This transformation function was then applied to transform mean fixation durations extracted from the published original studies into the mode. In a final step, sampling period was determined as mode fixation duration plus mode saccade duration (as estimated from study 1) and then converted into a frequency value.

In brief (for details see Supplementary Methods), the development of this function involved (i) fitting ex-Gaussian distributions to the empirical distributions of fixation durations in 29 original datasets to which we had access (and which were not included into the meta-analysis), and (ii) retrieving distributional parameters for each fitted distribution (specifically: μ, the mean of the normally distributed component; σ, its standard deviation; τ, the parameter reflecting the rightward skew, representing the contribution of the exponential distribution). This allowed us to (iii) apply a regression analysis to predict the mode (dependent variable) from the mean fixation duration (predictor variable). Figure 6a presents the final generalized mean-to-mode transformation function, applied to all possible fixation durations in the range covered by the meta-analysis. Figure 6b shows how well the modes of our 29 datasets can be recovered by this function: Despite some unsystematic noise, the numeric transformation was nearly perfect (i.e., beta = .95; SE = .23; *t*(28) = 4.1). We then used the transformation function to estimate the respective modes from the 1,420 mean fixation durations of the meta-analysis dataset. To obtain the sampling period *t* (i.e., the interval from the onset of a saccade until the end of the following fixation; see also Study 1, above), a saccade duration estimate of 29ms (i.e., the mode of saccade durations from the reading dataset used in the first experiment) was added to each of the mode fixation duration. This is feasible since saccade durations do not differ much between persons during reading (see, e.g., Ref.^60^: range 20-39ms, mean: 29ms). Finally, the sampling period values (*t*) were transformed into frequency values (*f* = 1 / *t*).

**Figure 6.**
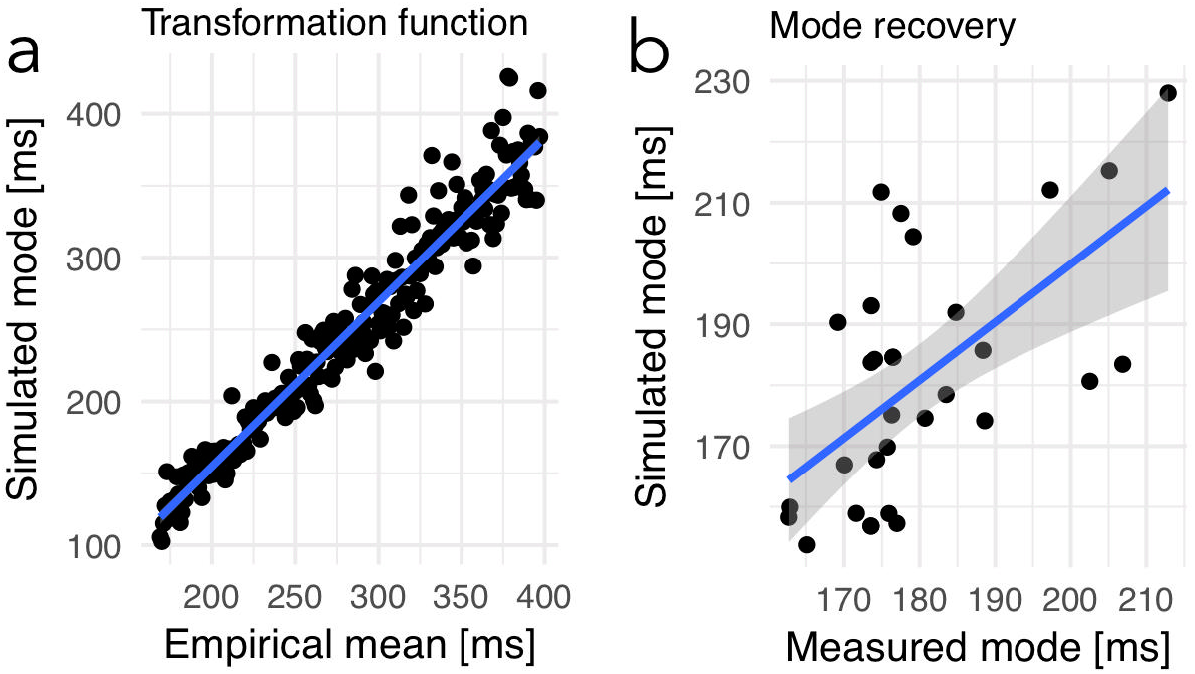
Transformation function for converting mean fixation durations into simulated mode values. The function was established using ex-Gaussian estimations on 29 empirical datasets containing fixation durations. For details see Supplementary Methods. (a) Performance of the final mean-to-mode transformation function (blue line) demonstrated here for all 248 possible fixation durations, i.e., for each millisecond within the range of fixation durations included in the meta-analysis (149 – 397 ms; x-axis: empirical means). (b) Performance of the final mean-to-mode transformation function, as demonstrated by the relationship between empirically measured (x-axis) and simulated (y-axis) modes from the 29 datasets used for establishing the transformation function.

### Writing system comparisons

In order to explore whether or not the sampling frequency of reading is influenced by global characteristics of writing systems and languages, we implemented four tests. First, reading Chinese (205 fixation duration data points) was compared to all alphabetic writing systems (1,215 fixation durations). Note that the Korean (with an alphabetic syllabary orthography^61^) and Japanese (using a mixture of Kana and Kanji) studies in the meta-analysis could not be clearly assigned to the character or alphabet categories and therefore were not included into this contrast. Second, among the alphabetic scripts we examined how the differences in transparency/opaqueness of the letter-to-sound relationship influence reading rates by a continuous predictor representing the number of grapheme-to-phoneme rules as defined by computationally implemented dual-route models^25^. A low number of grapheme-to-phoneme rules reflects a high transparency, meaning that letters more consistently represent only one speech sound. For example Italian, Dutch, and German are considered transparent orthographies, with 59, 104, and 130 rules, respectively ^25^). English and French, in contrast, are typically considered as in-transparent (opaque) with 226 and 340 rules, respectively, because letters map to multiple speech sounds on a regular basis. Third, we investigated the cross-linguistic relationship between variance in the peak modulation spectra from speech and mean sampling frequencies in reading. To this end, we retrieved the modulation spectra for Chinese, Dutch, English, French, Japanese, and Spanish from Figure 3c in Ref.^5^ and from Figure 7 in Ref.^6^. The modulation spectra varied from 4.3 Hz in English to 5.4 Hz in Dutch. Finally, in the fourth test, we investigated the relationship between eye movement sampling in reading and the information density of a language. This parameter indicates how dense a language codes meaning in texts^27^. The density is coded from 0 to 1 and could be retrieved for a subgroup of languages in the present meta-analysis dataset (i.e., Chinese, English, French, German, Italian, Japanese, and Spanish) from Ref.^27^. Density varied from dense languages like Chinese (0.94) to less dense languages like Japanese (0.49).

All four effects were analyzed using linear mixed models (LMM^62^). In addition to the parameters of interest, we accounted for experimental settings (experiment vs. corpus-based studies), for the different eye trackers used (which may also imply use of different saccade detection algorithms), and for different fixation measures reported (mean, single, or first fixation duration) by introducing these parameters into the LMMs as fixed effects. Also, for the modulation spectrum and information density comparisons we added a factor contrasting character-based (i.e., Chinese) vs. alphabetic writing systems, to account for perceptual difficulties, and for all four LMMs we estimated the random effect on the intercept of study, to take into account unspecific differences between studies. t-values larger than 2 were interpreted as significant^63^.

## Acknowledgments

The research leading to these results has received funding from the European Community’s Seventh Framework Programme (FP7/2013) under grant agreement n° 617891 awarded to CJF and from the European Community’s Horizon 2020 Programme under grant agreement n° 707932 awarded to BG. The funders had no role in study design, data collection and analysis, decision to publish or preparation of the manuscript. We thank Jutta Müller for helpful comments on a previous version of the manuscript.

## Author contributions

B.G., D.P. and C.J.F. designed research; B.G. and S.H. performed Study 1; B.G. and J.G. performed Study 2 (Meta-analysis); B.G. and K.G. performed Study 3 and 4; B.G., A. T. and J.S. analyzed data; and B.G. and C.J.F. wrote the paper. All authors gave comments on the paper during the process.

## Competing interests

The authors declare no competing interests.

## Supplementary Methods: Mean to mode transformation function

The typical distribution of eye fixation duration data in reading is ex-Gaussian (see Figure 1c). For simplification, one can decompose the ex-Gaussian distribution in a normal and an exponential distribution. This decomposition is a simplification on a mathematical level since both the normal and exponential distributions can be modeled easily. Consequently, one can describe the central tendency of the ex-Gaussian distribution by three parameters: the mean and standard deviation of the normal distributed component and the exponential component (i.e., reflecting the skew of the ex-Gaussian distribution). The μ relates to the mean of the normal distribution. The σ refers to the standard deviation of the normal distribution. The τ describes the rightward skew, i.e., representing the contribution of the exponential distribution.

In a frequency analysis one investigates if a reoccurring event, in our case a saccade, has a temporal structure. In Experiment 1, we showed that the mode of the ex-Gaussian distribution of the sampling durations (fixation plus saccade duration) indicates the most common sampling duration, which was found to be the adequate metric for the frequency estimation (i.e., by showing comparable frequency estimates based on mode fixation duration and power spectral estimation approaches but not when using the mean fixation duration). In the eye-tracking literature on reading, it is more typical to report mean fixation durations. Reporting mean not mode fixation durations is a central problem of the current meta-analysis. Accordingly, we developed the mean-to-mode transformation function described here. With this function, we transform the mean fixation durations extracted from papers into mode values.

We implemented the mean-to-mode transformation function in three steps: (i) We fit ex-Gaussian distributions (i.e., by decomposition methods) to existing empirical datasets. (ii) We used fitted ex-Gaussian parameters (μ: mean of the normal distribution; σ: standard deviation of the normal distribution; τ: exponential component describing the rightward skew) to simulate new, informed, ex-Gaussian distributions to derive a transformation function. (iii) We optimize the transformation function to increase transformation accuracy.

### (i) Ex-Gaussian fitting to existing empirical datasets

First, we fitted the three ex-Gaussian parameters to 29 empirical datasets containing fixation durations (11 published studies, i.e., three German studies from our lab, (1–3), and multiple English studies (4–10) for which datasets were openly available) using the *mexgauss* function from the *retimes* package in *R* (11). Figure S1a shows two empirical and the respective simulated distribution, including the mean and mode of the distribution, exemplarily. Henceforth, we jointly refer to these 29 datasets as the ‘simulation data’. Combined we now obtained the exact mean and mode of 29 datasets as well as the μ, σ, τ for each dataset. Note, the main selection criteria for the datasets used in the present study was availability and accessibility of the raw fixation duration values.

### (ii) Using fitted ex-Gaussian parameters to simulate new informed ex-Gaussian distributions

With the fitted ex-Gaussian parameters, we estimated, in a next step, three robust linear regression models (rlm function in R from the MASS package; (12)). One for each of the ex-Gaussian parameters (μ, σ, τ) to predict the mean fixation duration (μ: 0.40, SE = 0.09, t = 4.6; σ: 0.14, SE = 0.09, t = 1.5; τ: 0.60, SE = 0.09, t = 6.7). Figure S1b shows the relationships of each parameter to the means from each study.

Now, we can simulate realistic ex-Gaussian distributions (with 500 samples) for any mean value with the *exGAUS* function from the *gamlss*.*dist* package in R (13). These distributions can be realized by the fitted linear regression coefficients (intercept, beta weight), which describe the relationship of each of the three ex-Gaussian parameter estimates to the mean of the dataset (see Figure S1b). For example, one can go to the graphics and see that with a mean fixation duration of 200 ms one can obtain a μ value of 140, a σ value of 30 and a τ value of 40. Having a value to each of the ex-Gaussian parameters, one can simulate an ex-Gaussian distribution. This simulation then allows us to estimate the mode of the distribution, in our case around 150 ms. As a consequence, one can directly relate the mode of 150 ms to the mean value of 200 ms.

To reduce estimation noise and increase robustness against outliers, we sampled ex-Gaussian distributions, not only for the 29 datasets available but for the whole range of mean fixation durations (149 and 397 ms) from the meta-analysis. From these 248 simulated ex-Gaussian distributions, we estimated the mode values relating each mean to a mode value. Finally, these related mean and mode values allowed us to generate a generalized mean-to-mode transformation function by only one linear regression (e.g., blue line in Fig. S1d). Note, to realize the function in a generalized way, we on purpose neglected the specific experimental manipulations of the different studies of our simulation data.

### (iii) Optimizing the transformation function

For initial quality control, we used the fitted linear regression coefficients (intercept, beta weight) from the transformation function to transform the mean fixation duration of each of the 29 simulation datasets into a simulated mode. Since we were also able to measure the mode of these datasets were able to compare the simulated to the measured modes for each dataset. In Figure S1c, Level 1, we present the residual errors of the 29 simulated modes, relative to the measured modes. The negative relationship between the measured mode and the residuals (i.e., simulated minus measured mode) indicated a systematic overestimation for low measured modes and underestimation for high measured modes. This is likely caused by imprecisions in the ex-Gaussian fitting procedure. To account for this systematic error, we corrected the simulated modes by a sequential procedure. First, we described the error by a linear model. This model is then used to predict the over/underestimation of a given mode. The prediction is then used to correct the simulate mode values. This correction procedure was applied two times.

Figure S1c, Level 3, shows that after this sequential correction procedure, the final transformation function does not include a systematic error that one would expect to be present in the residuals. Figure S1d, accordingly, shows the final, i.e., corrected, transformation-function, for all possible 248 mean fixation durations. Figure S1e shows the final quality check from the simulation dataset showing the relationship of the measured and simulated modes. Despite some unsystematic noise, this optimized transformation function showed a near-perfect beta of 0.95 (SE = 0.23; t = 4.1).

**Figure S1.**
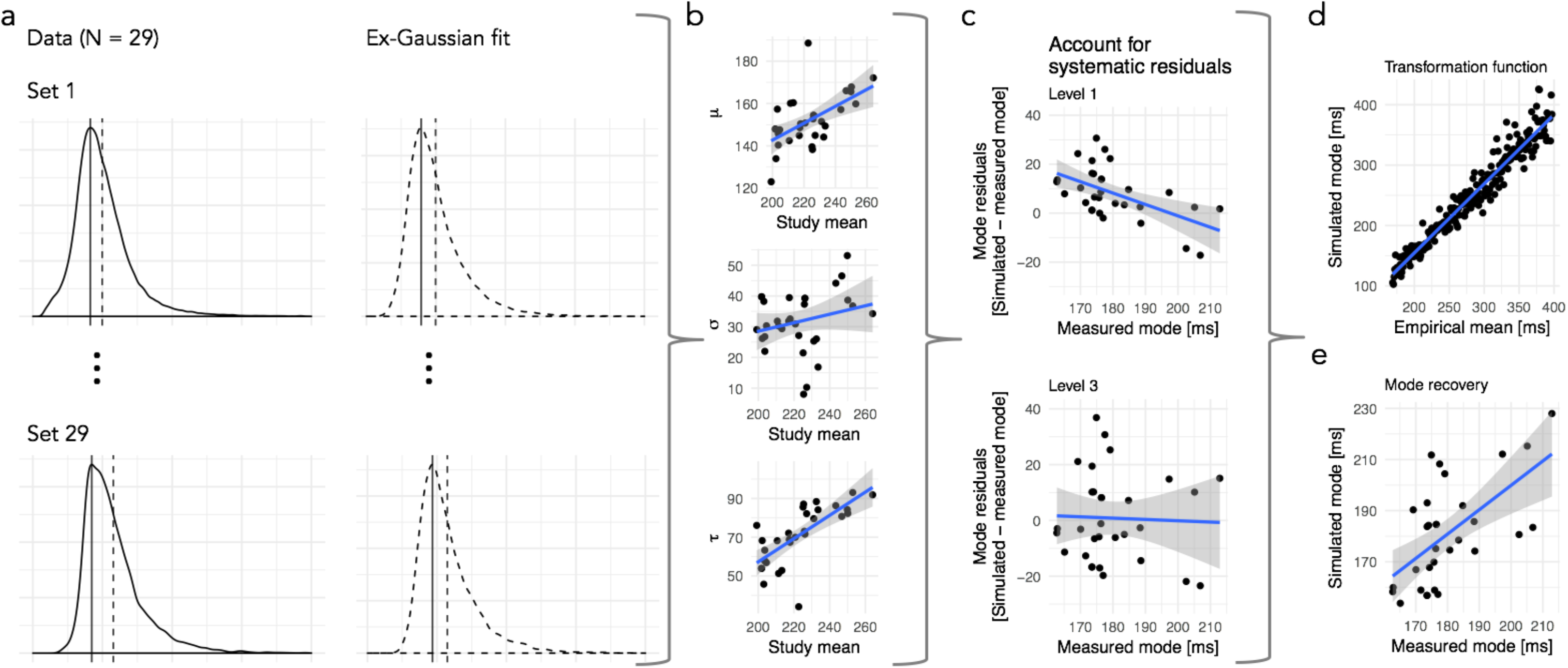
Development of a transformation function for converting mean fixation durations into simulated mode values. In order to estimate the predominant, i.e., mode, fixation duration from a single mean value extracted from a published empirical study, we implemented the following procedure: (a) First, an ex-Gaussian function was fitted separately to empirical fixation duration distributions of each of 29 independent datasets (left panel; see Materials and Methods), and we extracted the parameters μ, σ, τ describing the fitted ex-Gaussian distribution (right panel). (b) Second, across these 29 datasets, the relationship between the ex-Gaussian parameters and the empirical study means were described by linear models, separately for each parameter. The intercepts and beta-weights resulting from these linear models, for each ex-Gauss parameter, were then used to simulate an ex-Gaussian for each of the 29 empirical mean fixation durations, so that we could compare the empirical and simulated ex-Gaussian distributions. (c) Residuals for the mode estimation showed a systematic error, i.e., an overestimation for low modes and underestimation for high modes (upper panel / Level 1). We accounted for this systematic error, sequentially, by two linear models describing the error; see lower panel / Level 3 for residuals after accounting for the estimation error. (d) Performance of the final mean-to-mode transformation function (blue line) demonstrated for 248 fixation durations, i.e., for each millisecond within the range of the meta-analysis (149 – 397 ms). (e) Performance of the final version of the mean-to-mode transformation function, as demonstrated by the relationship between empirically measured and simulated modes from the 29 datasets used for establishing the transformation function.

## Supplementary Methods: Small talk interview

The full list of 22 questions from the “small talk” interview we conducted in German.

1. Wie viele Prüfungen hast du jetzt im Semester?
2. Warum studierst du? (ich frage meistens auch noch so allgemeiner, was genau sie studieren, wieso sie sich das rausgesucht haben, was ihnen daran Spaß macht)
3. Hast du dein Wohnort durch das Studium gewechselt?
4. Wie ist die Wohnungssituation für dich in Frankfurt?
5. Hast du schon früher eine Ausbildung/Studium gemacht?
6. Hast du schon mal ein Auslandsaufenthalt gemacht?
7. Hast du eine Zweitsprache? Welche Sprachen sprichst du?
8. Machst du irgendein Sport?
9. Spielst du irgendwelche Computerspiele / hast früher gespielt?
10. Was sind deine Hobbys / Interessen?
11. Was hast du am Wochenende gemacht? (wenn sie sich nicht erinnern können frage ich was sie für das kommende Wochenende vorhaben)
12. In welchen Ländern warst du schon?
13. Was ist dein Lieblingsessen? Was isst du gerne?
14. Wie findest du das Wetter im Moment so?
15. Was ist deine Lieblingsjahreszeit?
16. Machst du irgendein Nebenjob?
17. Hast du für den Sommer / die Weihnachtsferien etwas vor?
18. Isst du gerne in der Mensa?
19. Wo kommst du eigentlich her?
20. Was machst du im Studium im Moment so inhaltlich?
21. Was hast du nach dem Studium vor?
22. Was war dein letzter Kinofilm / Fernsehfilm / Serie, die du geguckt hast?

## Notes

### Competing Interest Statement

The authors have declared no competing interest.

### Summary of Updates

The main change to the previous version is the addition of Study 4, including a new, pre-registered, empirical investigation directly comparing the eye movement sampling rate and the speech rate in native German speakers.

